# Enhancing protein structure prediction accuracy by prioritizing important residues using protein language models

**DOI:** 10.1101/2025.09.28.679101

**Authors:** Yu Liu, Boming Kang, Qinghua Cui

**Author notes:** To whom the correspondence should be addressed: Qinghua Cui.

## Abstract

Accurate prediction of protein tertiary structures from amino acid sequences remains a fundamental challenge in computational biology. Although AlphaFold2 represents a major advance, systematic discrepancies persist between its predictions and experimentally determined structures. Given that individual residues contribute differentially to protein function, we hypothesized that incorporating residue-specific importance metrics could improve prediction accuracy. Here, we develop *i*-Fold (*importance*Fold), an enhanced neural architecture enhances the AlphaFold2 architecture by integrating protein language model ESM-derived residue importance scores (RIS) as dynamic positional weights during training. Our approach dynamically weights amino acids using RIS during structure prediction, thereby directing computational attention toward functionally critical residues and regions. Evaluation on a benchmark test set of 3,559 protein structures reveals that *i*-Fold significantly improves accuracy (reduction in r.m.s.d., *p* = 0) and achieves a higher prediction success rate (7.6% improvement: 55.1% → 62.7%). Notably, *i*-Fold demonstrates particular improvements for targets that are typically challenging for AlphaFold2, including ribosomal proteins, membrane proteins, and orphan proteins. Consistent results were obtained on a completely independent test set of 167 recently released protein structures, where *i*-Fold again exhibited a higher prediction success rate (6.0% improvement: 43.7% → 49.7%) compared to AlphaFold2. Our findings indicate that explicit integration of RIS can advance the state-of-the-art in protein structure prediction, producing more accurate and generalizable models without substantially increasing computational cost.

## Introduction

Protein three-dimensional structure determination is a cornerstone of structural biology, as the intricate relationship between form and function governs the behavior of biological macromolecules. Precise structural insights enable a wide range of applications, from rational drug design^[1]^ and enzyme engineering^[2]^ to mechanistic understanding of disease pathology^[3]^. While conventional techniques such as X-ray crystallography and cryo-electron microscopy deliver atomic-level accuracy, they still face challenges in throughput and resolution^[4]^. These limitations have spurred the rapid development of artificial intelligence (AI) for protein structure prediction, where deep learning algorithms have demonstrated remarkable progress in deciphering the folding patterns of amino acid sequences.

AlphaFold, the AI tool developed by Google DeepMind, has marked a watershed moment in this field. Its successor, AlphaFold2, gets a further upgrade^[5]^. By integrating attention mechanisms and end-to-end geometric learning, this deep learning system achieved unprecedented backbone accuracy of 0.96 Å in the CASP14 competition, rivaling experimental methods^[6]^. Subsequently, several adaptations of the AlphaFold algorithm emerged, such as ColabFold^[7]^ and OpenFold^[8]^, which have expanded its accessibility and adaptability. Parallel developments in protein language models, particularly ESM^[9]^, have introduced complementary approaches through self-supervised learning on evolutionary sequences to capture complex residue-residue dependencies. These models excel at predicting mutation effects and functional sites without requiring explicit structural information^[10]^, thereby offering both an alternative to labor-intensive multiple sequence alignment (MSA) and novel perspectives on protein structure prediction^[11, 12]^.

Despite these advances, critical limitations persist in current methodologies, necessitating further refinement^[5, 13-15]^. AlphaFold demonstrates reduced accuracy in predicting multi-domain proteins and dynamic conformational changes^[16, 17]^, while its heavy dependence on MSA limits its applicability to evolutionarily isolated sequences^[18]^. In contrast, ESM-based approaches, though independence of MSA and computationally efficient, demonstrate lower precision in three-dimensional coordinate prediction compared to AlphaFold^[10, 19]^. These constraints highlight the urgent need for innovative solutions that bridging existing gaps in accuracy, versatility, and applicability.

Position-based weighting has emerged as a powerful paradigm in biological sequence analysis. In RNA secondary structure prediction, for instance, models incorporating SHAPE reactivity as positional constraints have proven significantly more accurate than sequence-only approaches^[20]^. Similarly, in protein engineering, ECNet—which integrates local residue co-evolution information as positional weighting—consistently outperforms unsupervised learning models and global representation methods^[21]^. In addition, we previously proposed computational methods for quantifying the importance of mRNAs/lncRNAs and single nucleotides, demonstrating their utility in predicting mutation effects, cancer prognosis, and the transmissibility and death risk of SARS-Cov-2^[22-25]^.

Building upon these foundations, here we present *i*-Fold (*importance*Fold), an advanced structure prediction framework that enhances the AlphaFold2 architecture by integrating ESM-derived residue importance scores (RIS) as dynamic positional weights during training. This innovative approach enables selective focus on functionally critical residues and regions, resulting in improved prediction accuracy across diverse test datasets compared to AlphaFold2. Our findings compellingly demonstrate the effectiveness of positionally weighted learning strategies for next-generation protein structure prediction.

## Materials and Methods

### Overview of the proposed method

Since individual residues within a protein sequence may contribute differentially to its function, we hypothesized that incorporating residue-specific importance metrics could enhance the accuracy of protein structure predictions. Unlike conventional approaches, our method assigns each amino acid in the protein sequence a weight based on its corresponding residue importance score (RIS), derived from the ESM protein language model (**Fig. 1**). The strategy dynamically prioritizes functionally critical residues and regions during structure prediction.

**Fig.1.**
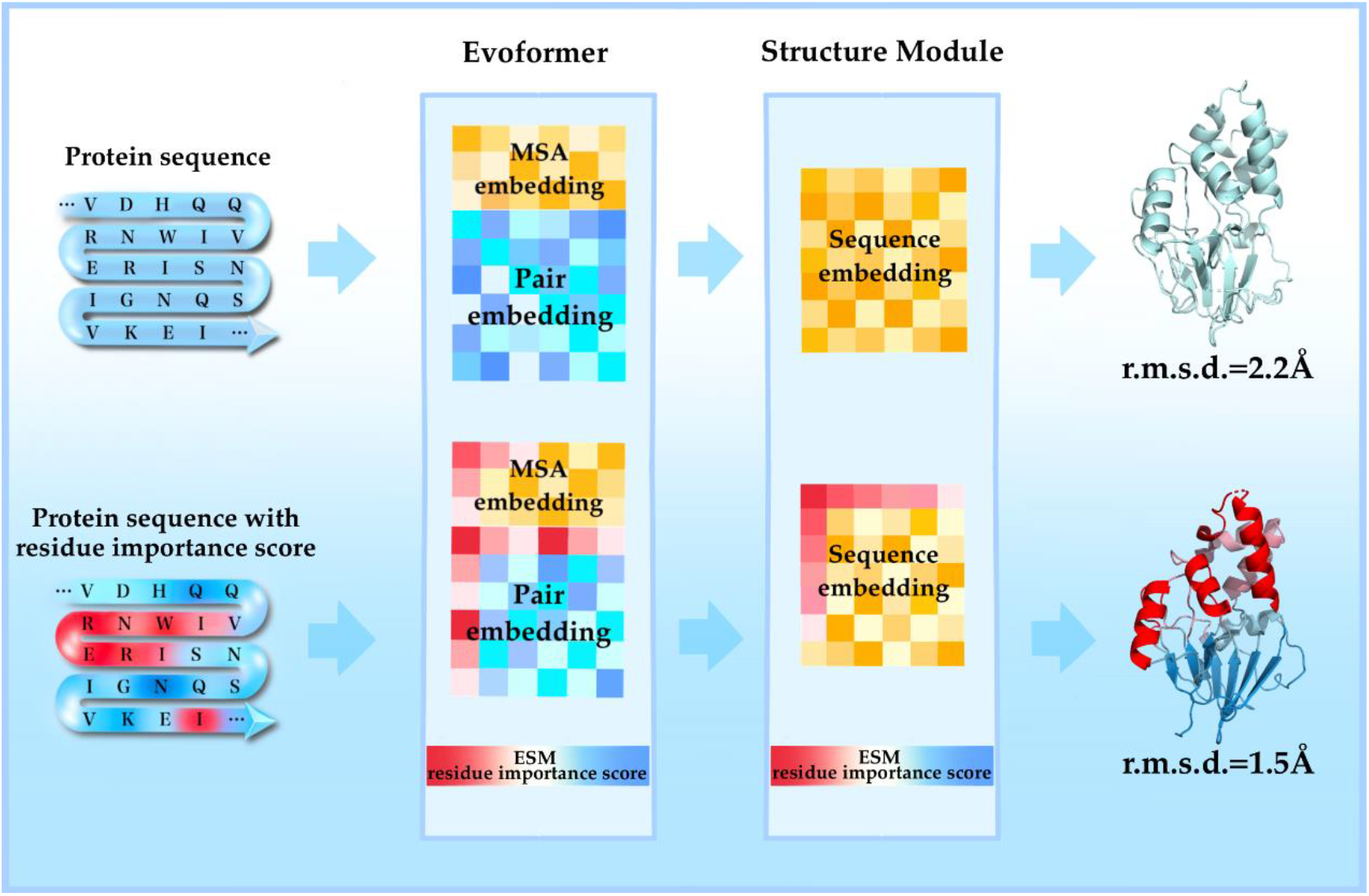
Overview of the proposed method, *i*-Fold (*importance*Fold). In contrast to the conventional strategy (upper panel, with all amino acids highlighted in light blue), the new approach assigns residue-specific importance weights—derived from the ESM large language model—to each amino acid based on its residue importance score (RIS). These weights are represented by distinct colors in the figure. By dynamically modulating the contribution of each amino acid during structure prediction according to its RIS, this method directs computational focus toward functionally critical residues and regions, enhancing the efficiency and biological relevance of the prediction process.

### Data preparation and preprocessing

All 185,158 protein structure files and their corresponding multiple sequence alignment (MSA) files—originally used for training AlphaFold2^[6]^ —were obtained from the Registry of Open Data on AWS (RODA). From these, a random sample of 3,599 sequences was selected to form the benchmark test set. The remaining sequences were processed following the methodology described by Gustaf et al.^[8]^, using the same sampling frequency as in AlphaFold2, to generate index and cache files. This produced a training dataset structured in the OpenFold DB format, compatible with the training script. Additionally, a completely independent test set comprising 167 protein structures published after June 1, 2024, was curated from the Protein Data Bank (PDB). MSAs for this set were generated using JackHMMER and HHBlits against the Big Fantastic Database (BFD) released with AlphaFold2.

### Model training

We used OpenFold^[8]^, a trainable PyTorch-based implementation of AlphaFold2, for all training tasks. Models were trained on four NVIDIA A100 GPUs using OpenFold’s script, with each run completing 10 over approximately seven days. *i*-Fold employed the same training parameters as AlphaFold2, except for the loss function. A fixed random seed was used throughout to ensure reproducibility.

### Residue importance-weighted loss function based on ESM

For each protein sequence, the protein language model ESM-2^[10]^ was used to generate an amino acid probability matrix *p* of dimensions (L, 20), where L is the sequence length. This matrix encapsulates evolutionary information per position. Information entropy was computed from *p* to quantify the mutation potential *E*_*i*_ at position *i* (Equation 1):

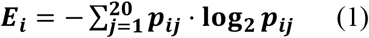

During evolution, conserved protein sites are more likely to form specific structural domains and perform key functional roles. Therefore, accurate prediction of structures in regions with lower mutation potential is particular important. Guided by this rationale, we developed residue importance score (*RIS*), which assigns higher weights to sites exhibiting lower mutation potential (Equation 2):

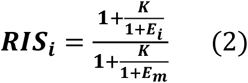

Here, *K* is a hyperparameter optimazed via grid search, and *E*_*m*_ is a constant representing the maximum value of the information entropy *E*. The final loss function was obtained by wighting the original FAPE loss with *RIS* (Equation 3):

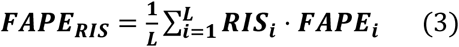

### Model performance evaluation

Root mean square deviation (r.m.s.d.) was used to evaluate structural similarity between predicted and experimental conformations. Calculations were performed using a custom script based on BioPython v1.84. In a limited number of protein cases where the .cif structure file and the input FASTA sequence file showed discrepancies, the following criteria were applied:

1. If the two sequences are of equal length and differed by fewer than three residues or contain non-standard amino acids, mismatched positions were ignored during alignment and r.m.s.d. calculation;
2. For sequences with terminal insertions or deletions, only the overlapping region is used for alignment and calculation;
3. Sequences with internal insertions or deletions, or those with an alignment length shorter than 10 residues, are excluded due to expected high structural divergence.

### Statistical analysis and visualization

All statistical analyses were performed using Scipy v1.7.0 and R v4.2.3. Results were visualized with the R package ggplot2, and protein structures were rendered using PyMOL.

## Results

### Development of a protein folding model incorporating residue importance score weighting

As shown in the diagram (**Fig. 1**), we hypothesized that incorporating residue-specific importance metrics could enhance the accuracy of protein structure predictions based on the observation that individual residues within a protein sequence may contribute differentially to its function. The new framework assigns each amino acid in the protein sequence a weight based on its corresponding residue importance score (RIS), derived from the ESM protein language model (**Fig. 1**).

AlphaFold2 utilizes the frame-aligned point error (FAPE) as its primary loss function during training^[6]^. For a given protein, prediction errors may occur either within functional domains or in linker regions (**Fig. 2A**). Under the conventional FAPE loss, both types errors contribute equally to the total loss. However, inaccuracies in functionally critical regions are more consequential^[26, 27]^ and should therefore incur a higher penalty. To address this, we computed the information entropy for each residue using the protein language model ESM-2, which generated a RIS. This RIS was then integrated into the loss function, yielding a modified version termed FAPE_RIS_.

**Fig.2.**
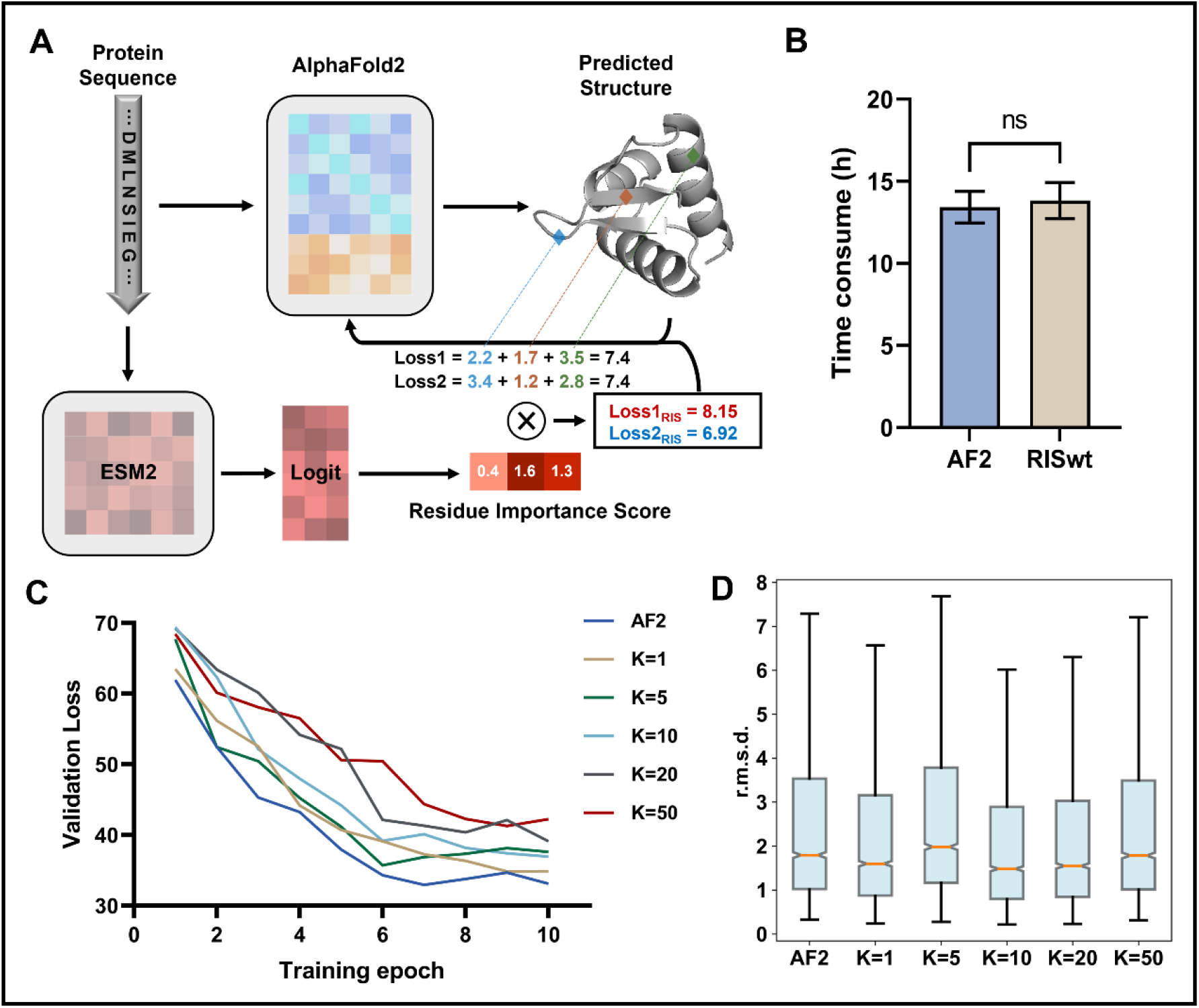
Development of a residue importance-weighted protein structure prediction model. **(A)** Schematic illustration of the loss calculation. Loss1 corresponds to a case where the prediction error is relatively high in functional domains, whereas Loss2 represents the case with a higher error in connecting regions. After applying RIS-based weighting, clear differences in loss values emerge between these two scenarios. **(B)** Comparison of training time per epoch. AF2: AlphaFold2; RISwt: RIS-weighted model. **(C)** Reduction in training loss across different values of the weight scaling factor *K*. **(D)** Comparison of prediction root mean square deviation (r.m.s.d.) across different values of the weight scaling factor *K*.

Retrained AlphaFold2 with FAPE_RIS_ did not substantially increase computational cost, as training times remained comparable (**Fig. 2B**). We explored various scaling coefficient *K* to optimize the weighting of ESM-derived RIS (see Methods). Training under different *K* values consistently reduced loss, demonstrating the stability of FAPE_RIS_ (**Fig. 2C**). Optimal performance was achieved at *K* = 10 (**Fig. 2D**), and this model was selected as the final version of *i*-Fold.

### *i*-Fold significantly enhances prediction accuracy

We compared *i*-Fold against AlphaFold2 on the benchmark test set of 3,599 experimentally resolved protein structures. The root mean square deviation (r.m.s.d.) between the predicted and experimental structures was used as the primary metric. As a result, *i*-Fold and AlphaFold2 r.m.s.d.s showed highly correlated r.m.s.d. (Spearman’s Rho=0.945, p = 0, **Fig.3A**, Supplementary File S1). Notably, *i*-Fold achieved significantly lower median r.m.s.d. (1.482 Å vs. 1.794 Å, *p* = 0, one-tailed paired Wilcoxon test, **Fig.3A**) than AlphaFold2, indicating globally improved accuracy.

**Fig.3.**
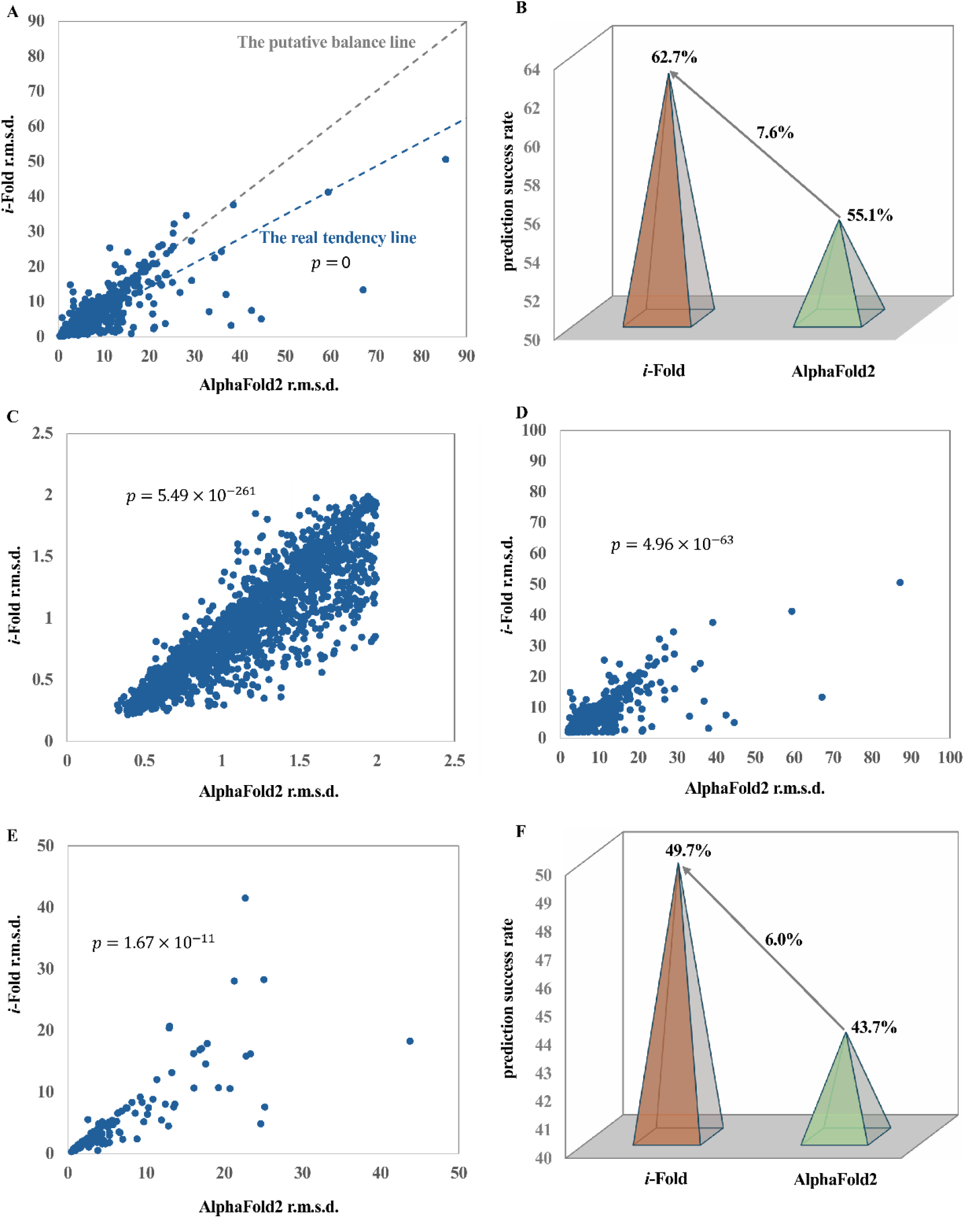
Performance comparison between *i*-Fold and AlphaFold2 on a benchmark test set (A-D) and a completely independent test set (E-F). **(A)** *i*-Fold achieved significantly lower median r.m.s.d. than AlphaFold2. **(B)** *i*-Fold also attained a higher prediction success rate compared to AlphaFold2. **(C)** On proteins correctly predicted by both methods, *i*-Fold yielded a lower median r.m.s.d. than AlphaFold2. **(D)** Similarly, for proteins that both methods failed to predict correctly, *i*-Fold still exhibited a lower median r.m.s.d. **(E)** On the independent test set, *i*-Fold again showed significantly lower median r.m.s.d. **(F)** *i*-Fold also achieved a higher prediction success rate in the independent test.

Using a common threshold of r.m.s.d. ≤ 2.0 Å to define successful predictions^[6, 8, 28]^, *i*-Fold attained a success rate of 62.7% (2257/3599), outperforming AlphaFold2 (55.1%, 1983/3599) by an absolute margin of 7.6% (*p* = 6.15 × 10^−11^, Two-Proportion Z-test, **Fig.3B**). Moreover, *i*-Fold also demonstrated superior accuracy both on proteins correctly predicted by both methods (n=1955; *p* = 5.49 × 10^−261^, **Fig.3C**, Supplementary File S2) and those both methods failed (n=1314; *p* = 4.96 × 10^−63^, **Fig.3D**, Supplementary File S3).

To further validate these findings, we assembled an independent test set of 167 recently published PDB protein structures (Supplementary File S4). *i*-Fold outperformed AlphaFold2 again, reducing the median r.m.s.d. from 2.61 Å to 2.03 Å (*p* = 1.67 × 10^−11^, one-tailed paired Wilcoxon signed-rank test, **Fig.3E**). It successfully predicted 49.7% (83/167) of these structures within 2.0 Å r.m.s.d., compared to 43.7% (73/167) for AlphaFold2 (**Fig.3F**).

Next, it is interesting to explore the biological annotations for the proteins on which *i*-Fold significantly outperformed AlphaFold2 (*i*-Fold-genes, r.m.s.d. <= -2.0 Å) and those AlphaFold2 significantly outperformed *i*-Fold (AlphaFold2-genes, r.m.s.d. >= 2.0 Å). For doing so, we performed gene ontology (GO) enrichment analysis using the WEAT tool^[24]^. While no terms were enriched among proteins better predicted by AlphaFold2, *i*-Fold-genes showed significant enrichment for terms related to translation, rRNA, ribosome, and membrane-associated biological process (BP), molecular function (MF), and cellular component (CC) (full list in Supplementary File S5, **Fig.4A-C**). This suggests that *i*-Fold is particular advantageous for predicting structures of proteins that are traditionally challenging to resolve.

**Fig. 4.**
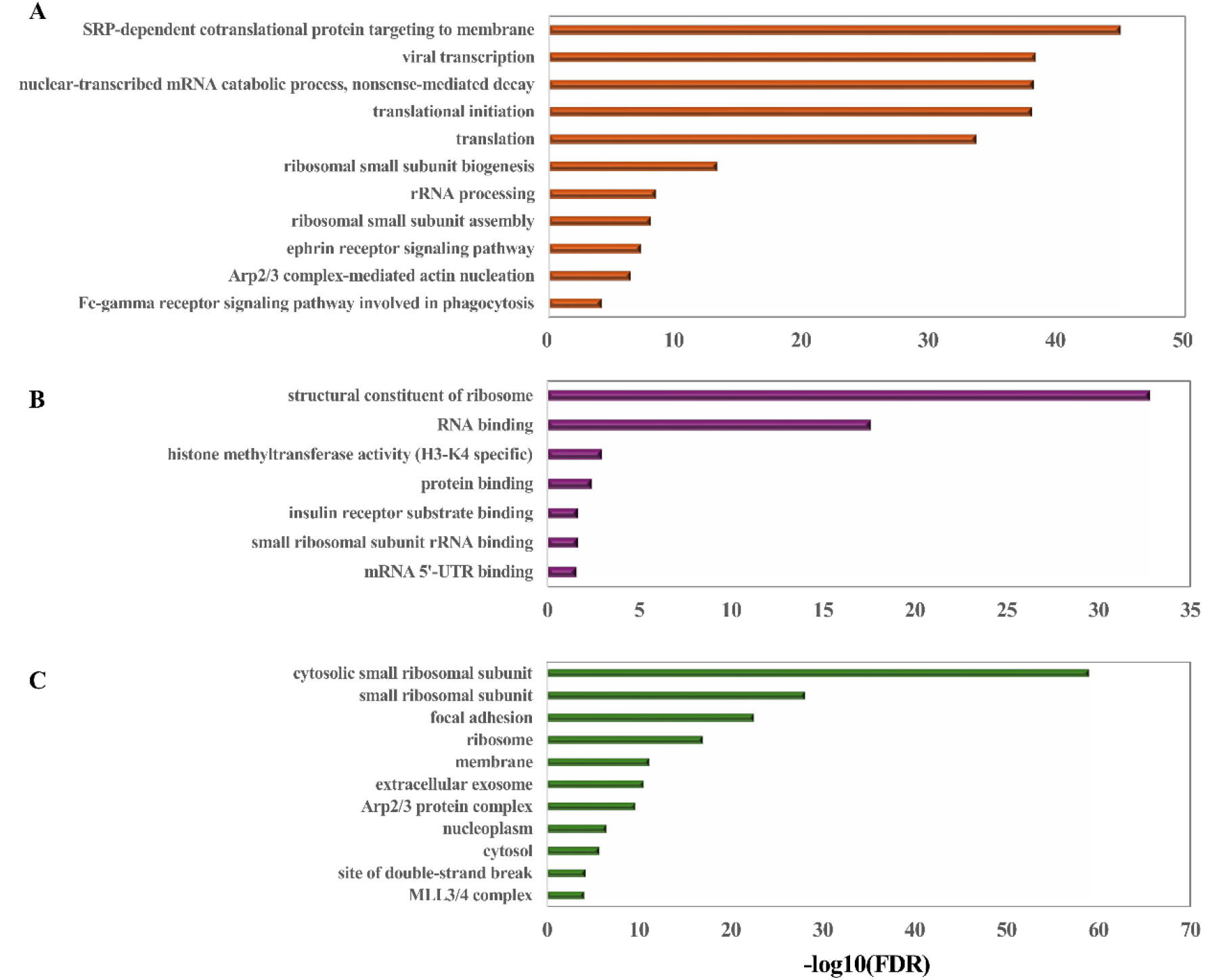
Enriched Gene Ontology (GO) terms for proteins in which *i*-Fold significantly outperformed AlphaFold2. **(A)** Biological process (BP). **(B)** Molecular function (MF). **(C)** Cellular component (CC).

### RIS improves protein structure prediction through multiple mechanisms

We further investigated how RIS enhances prediction accuracy. AlphaFold2’s performance is known to depend heavily on the quality of multiple sequence alignments (MSAs), often resulting in poor predictions for low-homology sequences^[29, 30]^. Indeed, on sequences with fewer than 500 MSA hits, AlphaFold2 yielded an average r.m.s.d. of 5.23 Å, whereas *i*-Fold reduced this by 1.08 Å (**Supplementary Fig. 1**). For example, the chloride channel protein (PDB: 7JNA), which retrieved only 60 homologous sequences during the MSA step^[31]^, was poorly modeled by AlphaFold2 at the N- and C-terminal α-helices. *i*-Fold produced a more accurate prediction (**Fig.5A**). Considering ESM’s ability to capture evolutionary information^[10]^, we hypothesize that the improved performance of *i*-Fold on low-homology sequences is primarily attributable to ESM’s capacity to partially compensate for the limitations of MSA by providing additional evolutionary context.

**Fig. 5.**
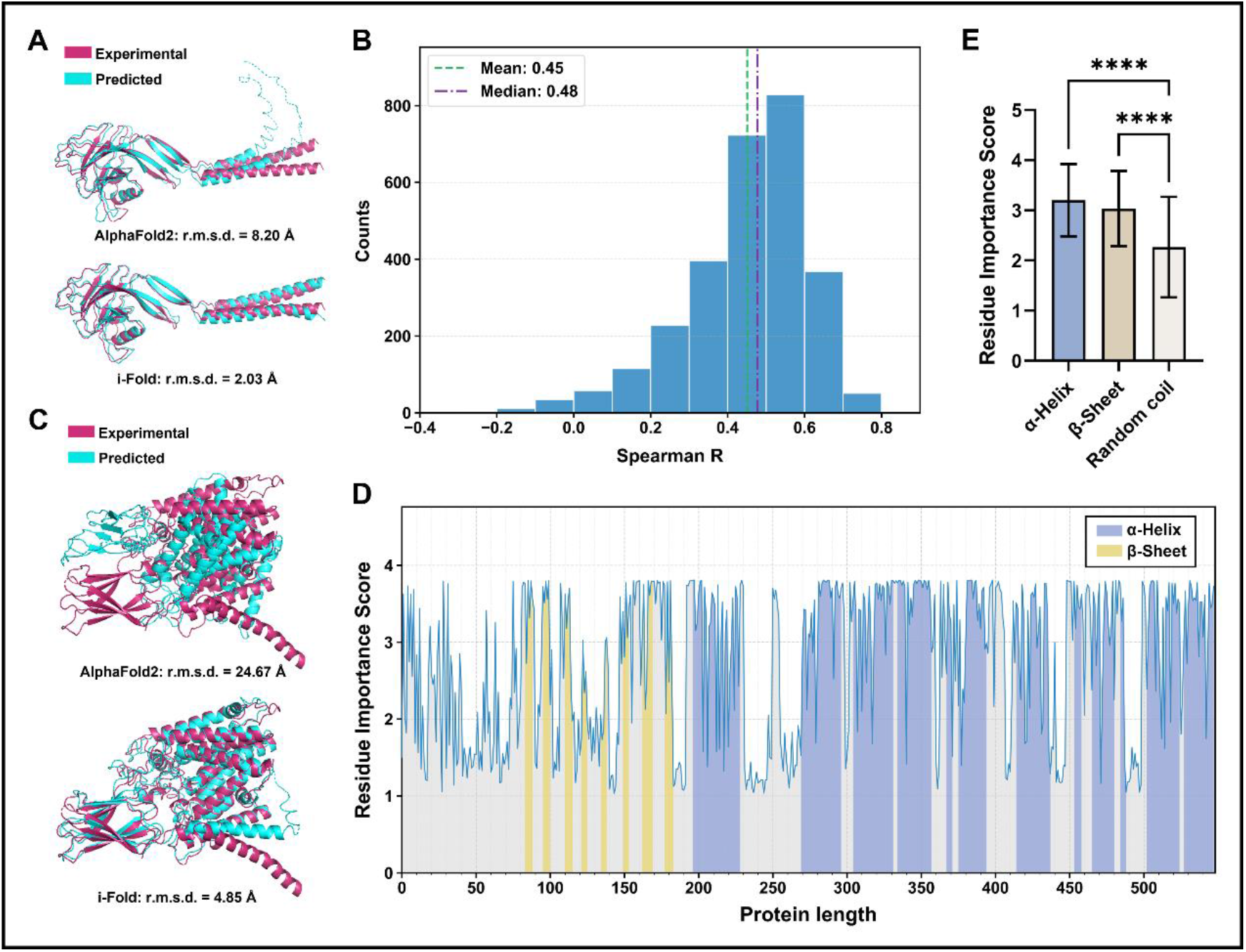
Mechanism underlying the improvement of prediction accuracy through residue importance scores. **(A)** Predicted structures of protein 7jna generated by AlphaFold2 and *i*-Fold. **(B)** Correlation between residue conservation and residue importance scores for sequences with MSA count > 1000 (n = 2889). **(C)** Predicted structures of protein 8vkj generated by AlphaFold2 and *i*-Fold. **(D)** Residue importance scores and corresponding secondary structure elements along the full-length sequence of 8vkj. **(E)** Comparison of importance scores among residues with different secondary structures, as shown in (**D**). Error bars indicate standard deviations; ****p < 0.0001.

Notably, *i*-Fold also improved predictions for high-homology sequences with. Among 2,889 proteins with >1000 MSA hits, ESM RIS weights correlated only moderately with MSA-derived conservation scores (Spearman’s Rho ≈ 0.45, **Fig. 5B**), indicating that RIS captures information beyond pure evolutionary conservation. For instance, in the lysosomal membrane protein HGSNAT (PDB: 8VKJ)^[32]^, AlphaFold2 failed to accurately model a β-sheet domain within the extracellular region, while *i*-Fold correctly predicted both domains (**Fig.5C**). Visualization of ESM RIS weights, we found a strong correspondence between regions with high RIS weights revealed strong agreement between high-weight regions and those forming specific secondary structures (**Fig. 5D-E**). In summary, ESM-derived RIS improves structure prediction through two complementary mechanisms: supplementing MSA-based evolutionary information and highlighting regions prone to forming specific structural elements.

## Discussion

In this study, we develop *i*-Fold (*importance*Fold), an advanced neural architecture that builds upon AlphaFold2’s framework while integrating residue importance scores (RIS) derived from the ESM protein language model. Evaluations across diverse datasets demonstrate that *i*-Fold achieves overall improved performance compared to AlphaFold2, with particularly notable gains in specific protein categories.

The statistically significant reduction in average r.m.s.d., along with the improved prediction success rate, strongly support our hypothesis that protein structural prediction benefits from residue-specific weighting. This finding aligns with emerging perspectives in computational biology that highlight the importance of functional hierarchies within protein structures^[33-35]^. Notably, *i*-Fold exhibits superior performance in predicting translation-related proteins and membrane structures—categories often rich evolutionarily conserved functional motifs. The residue importance weighting mechanism effectively directs computational attention to critical interaction interfaces, thereby improving predictive accuracy in these categories. Furthermore, *i*-Fold demonstrates robust performance on orphan proteins—those with limited homologous sequence information, which are typically challenging for AlphaFold2. This improvement suggests that ESM-derived residue importance weights can effectively compensate for sparse evolutionary signals in multiple sequence alignments (MSAs). Unlike methods that directly integrate ESM embeddings as additional inputs^[10, 36]^, our approach consistently enhances performance without introducing new parameters, while also offering improved interpretability through explicit residue importance visualization. Given the quadratic scaling of attention mechanisms with sequence length, this parameter-efficient strategy is computationally advantageous.

Beyond benchmark evaluation, *i*-Fold maintained its performance improvement on a completely independent and newly released dataset, indicating that the residue importance metric captures intrinsic sequence-structure relationships rather than dataset-specific artifacts. This level of generalizability addresses a key limitation in previous attempts to incorporate auxiliary biological information, such as the contact-map strategy used in trRosetta^[37, 38]^.

Several mechanisms may explain for these improvements. First, ESM-derived residue importance scores effectively identify evolutionarily constrained residues that are critical for forming complex structural domains. Accurate placement of these critical residues simplifies the modeling of connecting regions. Second, while conventional approaches, using either MSAs or ESM embeddings, rely on attention mechanisms to emphasize inter-residue dependencies, they often underrepresent residue-specific features. In contrast, our method maps the evolutionary information captured by ESM directly onto individual residues through a weighted loss function, thereby mitigating inherent limitations of traditional methods.

Looking ahead, *i*-Fold can be further refined in several directions. First, although evolutionary conservation serves as a well-validated importance metric—as demonstrated across diverse protein classes—recent studies suggest that additional physicochemical factors also influence structural stability, such as hydrophobic gradients and electrostatic complementarity^[39]^. Future developments could therefore benefit from a multi-dimensional importance scoring system that integrates evolutionary, energetic, and dynamic constraints in a synergistic manner. This highlights the need for more comprehensive and refined approaches to evaluating residue importance, which represents a promising direction for future research. Second, while this study focuses on single-chain protein prediction, the growing importance of biomolecular complexes in structural biology^[28]^ motivates extending the importance-weighted framework to protein-protein and protein-ligand interactions. Third, although *i*-Fold is developed upon AlphaFold2, the concept of residue importance weighting could be adapted to enhance other AI-based protein structure prediction tools.

## CODE AVAILABILITY

All source codes of *i*-Fold algorithm have been deposited at https://github.com/LiuYu-pku/i-Fold.

## ACKNOWLEDGEMENTS

This study was supported by the grants from the National Natural Science Foundation of China [62025102].

## CONTRIBUTIONS

Q.C. proposed the original idea and supervised the study. Y.L. performed the study. B.K. provided helps in implementing the algorithm. Y.L. and Q.C. wrote the manuscript.

## Figure Legends

**Supplementary Fig. 1.**
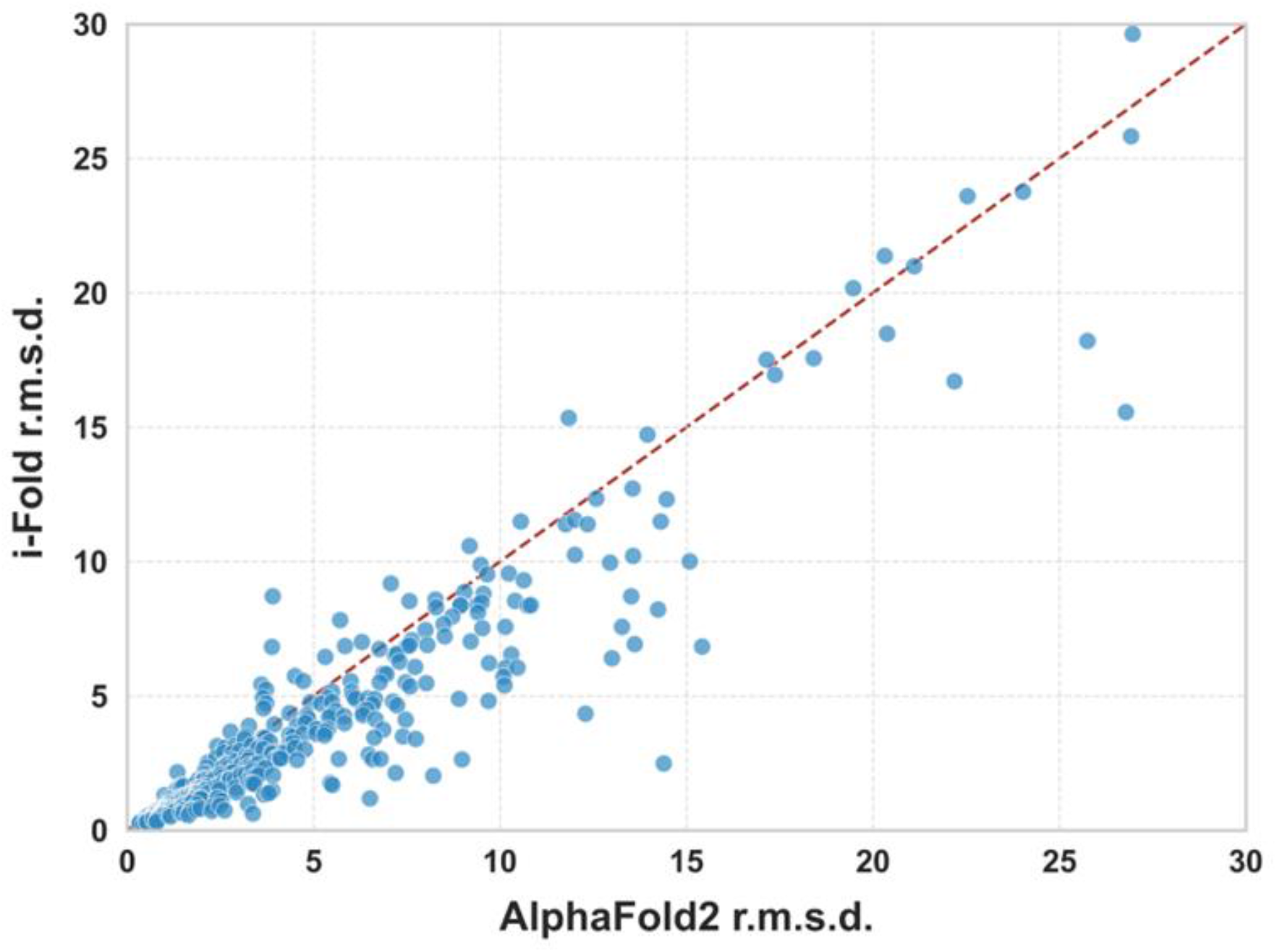
Comparison of the prediction performance between AlphaFold2 and *i*-Fold on low-homology proteins (n = 400) with fewer than 500 MSA hits.

